# Multifractality distinguishes reactive from proactive cascades in postural control

**DOI:** 10.1101/2020.10.21.349589

**Authors:** Damian G. Kelty-Stephen, Mariusz P. Furmanek, Madhur Mangalam

**Affiliations:** Department of Psychology, Grinnell College, Grinnell, IA 50112, USA; Department of Physical Therapy, Movement and Rehabilitation Sciences, Northeastern University, Boston, MA 02115, USA; Institute of Sport Sciences, The Jerzy Kukuczka Academy of Physical Education, 40-065 Katowice, Poland

**Author notes:** E-mail addresses:* (D. G. Kelty-Stephen), (M. Mangalam). These authors contributed equally to this work (Damian G. Kelty-Stephen, Madhur Mangalam).

**Keywords:** center of pressure, heavy tail, multifractality, quiet stance, multiscale PDF, scale invariance

## Abstract

Intermittency is a flexible control process entailing context-sensitive engagement with task constraints. The present work aims to situate the intermittency of dexterous behavior explicitly in multifractal modeling for non-Gaussian cascade processes. Multiscale probability density function (PDF) analysis of the center of pressure (CoP) fluctuations during quiet upright standing yields non-Gaussianity parameters lambda exhibiting task-sensitive curvilinear relationships with timescale. The present reanalysis aims for a finer-grained accounting of how non-Gaussian cascade processes might align with known, separable postural processes. It uses parallel decomposition of non-Gaussianity lambda-vs.-timescale and CoP. Orthogonal polynomials decompose lambda curvilinearity, and rambling-trembling analysis decomposes CoP into relatively more intentional rambling (displacement to new equilibrium points) and less intentional trembling sway (deviations around new equilibrium points). Modeling orthogonal polynomials of non-Gaussianity’s lambda-vs.-timescale relationship allows us to differentiate linear from quadratic decay, each of which indicates scale-invariant and scale-dependent cascades, respectively. We tested whether scale-dependent and scale-invariant cascades serve different roles, that is, responding to destabilizing task demands and supporting the proactive movement to a new equilibrium point, respectively. We also tested whether these cascades appear more clearly in rambling rather than trembling sway. More generally, we test whether multifractal nonlinear correlations supports this capacity of postural control to this two-step differentiation: both into rambling vs. trembling, then into scale-dependent vs. scale-invariant cascades within rambling sway. The results supported these hypotheses. Thus, the present work aligns specific aspects of task setting with aspects of cascade dynamics and confirms multifractal foundations of the organism-task relationship.

## 1. Introduction

### 1.1. Intermittency in dexterous behavior

Intermittency is the capacity of a system to hover between multiple modes of behavior. This flexibility is a hallmark of dexterous behavior. Rather than doggedly locking itself into a task’s constraints, dexterous behavior respects a degree of looseness from those constraints. Dexterous behavior necessarily meets tasks demands to accomplish a task, but the dexterously behaving system remains open to events happening in the surrounding context just beyond those task constraints. Hence, intermittency appears in the flexibility with which movement systems meet task constraints but check in with the broader context, switching between the two—irregularly and as needed.

Intermittency is important because any given task usually unfolds within a focal space that fails to exhaust all the scales at which an organism can act and respond to context. Task completion requires that organisms constrain their behavioral degrees of freedom to this focal task space’s confines, but events can also unfold at finer and coarser grains that prompt new actions not immediately aimed at the task. We begin with a non-postural example to illustrate intermittency before applying these concepts to posture for the present investigation. For instance, the task of reading these words and making sense of them requires steering eyes across the text. However, your behavior may include and may even benefit from deviations: you may find that the ambient lighting needs adjusting, or you may get up to refill your coffee cup. Moreover, even then, when eyes are glued to the screen, the gaze exhibits fixational movements at specific points of focus and saccades to another part of the text. That is, even when they are entirely on task, organisms roam through the task and explore in an odd and idiosyncratic way, for example, scrolling skimming back and forth to better grasp the needed meaning. Organisms thrive on the ability to complete tasks, but they thrive as well on the ability to detach from the task constraints and attend to new events or details that may arise.

Postural control exhibits intermittency as well. Whereas eyes scan across the screen to search for a word, across a kitchen for coffee, and back to the screen, postural sway of an upright body spreads across a base of support in an ongoing search for stability. Much like eye gaze shows fixational movements around focal points or saccades from point to point, sway exhibits a similar sort of ability to perch in one equilibrium point and then to move to another. In between these saccade-like shifts in equilibrium points lie fixational-like movements as the postural system explores the full range permitted by the current equilibrium point. Previous work examining the postural center of pressure (CoP) has shown that this full range of sway exhibits intermittency and exhibits a stronger signature of intermittency under more destabilizing task constraints [1]. In this investigation we test whether the intermittency crucial for maintaining upright stance depends on the relatively more intentional aspect of postural sway, namely ‘rambling’ sway as distinct from the smaller and more directionless ‘trembling’ sway [2,3]. Specifically, we expect that rambling sway’s intermittency might reveal further detail about separate classes of cascades (i.e., separate classes of interactions across timescales), responsible for reacting to perturbation and proactively redirecting posture to a new equilibrium point.

The analysis of intermittency has often been a double-edged sword for movement science. As a hallmark of dexterity, intermittency warrants explicit acknowledgment in models of dexterous behavior. However, intermittency poses a major challenge to modeling itself. For instance, modeling requires that we address causal relationships in distinct symbolic terms that become amenable to mathematical description [4]. Indeed, some modeling approaches will understand this requirement as logically related to all modeled causal relationships having their source in independent causal factors [5]. Certainly, if we take the premise that all causal factors in the modeled system act independently, then the most elegant models will have clean one-to-one correspondences between causal factors and symbolic variables in the model. For this purpose, the Gaussian probability distribution function (PDF) becomes a central theme for all statistical attempts to project such models into the data. The Gaussian PDF is the logically entailed, summed consequence of very many independent random variables, leading us to expect behaviors to exhibit Gaussian distributions centered on the system’s independent contributions of causal constituents. And where our independence-assuming statistical model does not fit the entirety of the observed behavior, this approach expects the residuals (measurement minus model) to exhibit homogeneity, specifically in terms of identical, independent distribution across space and time [6].

However, intermittency throws a wrench into this modeling strategy for at least one of two reasons. The first reason is easy to brush off as oddity of the measurement: intermittency guarantees heterogeneity, and so fitting a Gaussianity-assuming model to the measurement of an intermittent behavior will leave residuals that fail to be identical or independent. From the premise that all causal constituents of the system are necessarily independent, we might tune the model to the data with lagged values of our measurement and introduce an unbounded set of new independent factors to our model [7]. This data-driven approach justifies its opportunism by a commitment to the independence of causal factors. However, the second reason intermittency raises challenges to modeling has to do with the possibility that this commitment to the independence of causal factors is erroneous. That is, intermittency may reflect the fact that causal constituents are primarily interactive and not independent.

Hence, whereas Gaussianity reflects homogeneously-behaving systems composed of independent factors, interaction-driven intermittency implicates non-Gaussianity. Indeed, we could make a schematic, conceptual analogy between intermittency and the bell-shaped curve of a Gaussian PDF: the middle portion could reflect relatively more frequent close compliance with the task constraints, and the sparser tails could reflect those perhaps-adaptive, −context-sensitive deviations that allow exploring a more distant relationship with task constraints. However, intermittency entails wider bells in the PDFs decaying in heavier tails than we find in a Gaussian PDF. Indeed, a familiar contrast from introductory-statistics training might be the difference between the Gaussian and the Student’s *t* distributions: the *t* has heavier tails that place more of the area under the curve to ensure that our inferential tests are more conservative when we do not know the population standard deviation.

Movement science thus has an important choice of how to study intermittency. It can persist with attempts to explain this important hallmark of dexterity with a modeling strategy at its Gaussian root, logically at odds with the predicted outcome. However, in this manuscript, we present an alternative modeling strategy that explicitly uses non-Gaussianity to model intermittent behavior. Specifically, the non-independence of constituent factors supporting intermittency entails at least two related commitments, one theoretical and the other operational and quantitative. The theoretical commitment is to cascade dynamics following from nonlinear interactions across multiple scales. Cascade dynamics entail multifractal structures, and so the multifractal formalism remains a compelling framework for modeling intermittency and explaining dexterous behavior. The operational and quantitative commitment is to the heavy-tailed form of lognormal distributions generated by nonlinear interactions across scales. The interest in examining PDF heavy tails for signs of nonlinear interactivity is not new to the study of goal-directed behavior [10,11], but this program of querying heavy tails has been fraught by the instability of inevitably small-sample tailed behavior and often by a contrived theoretical organization of alternative distributions between ‘heavy-tailed’ and Gaussian [12–14]. So far, these attempts have mostly proceeded independently of explicit mooring in multifractal evidence of cascade-like nonlinear interactions across scales—indeed, it has been long recognized that heavy tails are not straightforwardly evidence of cascades [15]. The present work aims to situate this investigation of intermittency on the firmer multifractal footing.

Subsequent remarks will be as follows. We first elaborate these theoretical and quantitative commitments. Then we introduce the multiscale method developed in hydrodynamics for estimating the non-Gaussianity of measured PDFs and outline how it uses the multifractal formalism’s strategy of *q* moments to accentuate the middle part of the PDF while de-emphasizing the tails. We then make the case that non-Gaussianity provides a continuous, multiscaled measure of intermittency untroubled by the tail fitting. Previous work has shown that postural sway in quiet standing exhibits non-Gaussian PDFs that depends on the interaction of destabilizing task demands with multifractal nonlinearity [1,16]. So, we outline hypotheses that further clarify how multifractal nonlinearity in sway supports the task-dependence of non-Gaussianity. These hypotheses will highlight how intermittency in postural sway brings into relief subcascades within the relatively more intentional ‘rambling’ equilibrium-setting sway and not in trembling sway. These subcascades will be specific either to the sensory reaction to destabilization or to proactive stabilizing movements. Lastly, we outline precisely why we expect these effects to be clearer in non-Gaussianity and more ambiguous in heavy tails.

### 1.2. Intermittency entails commitments to cascade dynamics and non-Gaussianity at multiple timescales

An important candidate theoretical reason for dexterity to manifest intermittency is cascade dynamics— the capacity for interactions across scales. A system operating simply on Gaussian terms can compartmentalize its behaviors cleanly towards task compliance—as cleanly as the experimenter aims to compartmentalize behavior between levels of a well-controlled manipulation. However, deviations found in practice reflect that the alternative point that dexterous behavior is rooted in structure and dynamics extending beyond the task constraints. For instance, we may only want that second cup of coffee while reading because we did not get enough sleep the night before. The task of ‘reading the manuscript’ is too narroq to entail or constrain whether we need to get that second cup of coffee. Non-Gaussianity of intermittency reflects cascade dynamics in which events at longer timescales ramify and reshape events at shorter timescales, and in which the latter events can percolate and coalesce into the former events. We suggest that dexterous behavior is fundamentally cascade-like, generating a fluid movement between task constraints and context-checking deviations beyond those constraints. Certainly, our theoretical investment in cascade dynamics comes with risks—fluidity brings the threat that dexterous behavior will sometimes slip and fall. But this investment in cascade dynamics promises immense returns to movement science that we could explain and model the flexible adaptivity long championed as central to dexterity.

The operational, quantitative commitment is to multifractal formalism’s expectation of non-Gaussian and specifically lognormal distributions. Just as the Gaussian distribution is the entailment of independent factors summing together, the lognormal distribution is the logical entailment of interactions amongst those factors. The lognormal reflects an understanding of ‘interaction’ literally as multiplications of random factors rather than their sum [10]. It indicates a distinct signature of intermittency, that is, a class of PDFs whose bell-like shape would grow tails heavy enough to be non-Gaussian. So, in one sense, heavy tails and non-Gaussianity go hand in hand, but although heavy tails are symptomatic of non-Gaussianity, the quality of non-Gaussianity is more central to intermittency than heavy tails are. Hence, there is more to non-Gaussianity than simply heavy tails, and the only reason to direct our attention to heavy tails is that they very often belong to whole distributions generated by interactivity sooner than by independence. However, the focus on heavy tails alone has risked analytically omitting the bulk of the measured behaviors. It is important not to forget that interactivity could well leave its statistical imprint on the measured behavior’s entirety. That is, interactivity is not only in play for extreme events. For this reason, Kiyono and colleagues [17–21] have developed a measure of non-Gaussianity that emphasizes variance, a statistic addressing the entirety of the distribution. Like multifractal scaling methods [22], their multiscale PDF analysis estimates variance after detrending at many scales. However, it respects that lognormal variance is defined differently than Gaussian variance. The measure of non-Gaussianity estimates the potentially non-zero lognormal variance that the observed PDF yields above and beyond the Gaussian variance.

What makes multiscale PDF a superior method for articulating cascade dynamics is its capacity to address the bulkier midsection of the PDF much more than the tails alone. The traditional maximum likelihood estimation (MLE) modeling is more sensitive to the PDF tails’ raw values. This sensitivity allows rare outliers to make otherwise light-tailed distributions (e.g., gamma or exponential) seem heavy-tailed like a lognormal distribution [23]. On the other hand, multiscale PDF analysis fits the more populated region between the sparsely-populated tails. It tests whether the bell-like shape is peaked in a way that is consistent with non-Gaussian distributions. Multiscale PDF accomplishes this emphasis on the middle portion of the PDF through its roots in the multifractal formalism. The multifractal formalisms use a *q* exponent to emphasize different-sized fluctuations—with higher and lower *q* emphasizing larger and smaller fluctuations, respectively [24]. Multiscale PDF uses a low value of q that spreads the midsection of the bell-shaped curve outward and retracts the tails. Accentuating the well-populated middle at the expense of poorly populated tails allows diagnosing non-Gaussianity even when tails are not heavy.

Multiscale PDF analysis thus amounts to a significant step forward in modeling intermittency, modeling PDF shape in a way that complements what multifractal analysis accomplishes in the time domain. The multiscale PDF method elegantly disposes of tail-focused questions that have long plagued this work, that is, over outliers [23], where the tail ‘starts’ [25], and how to make theoretical sense of piecewise/composite/mixture models presuming to reattach the tail to light-tailed distributions [26–28]. It grounds the discussion of intermittency on the full distribution and on how it may generate more or less lognormal variance at various scales. The non-Gaussianity profile across scales allows a more subtle view in which we can discern different cascade processes within the same measured behavior. We hope in this manuscript to bring this exquisite lens to the cascade dynamics in human postural control, and we aim to show that multiscale PDF analysis allows us to be specific about precisely which cascade processes support which aspects of dexterous behavior.

### 1.3. Hypotheses regarding the reactive and proactive cascades composing postural control in rambling sway

Our previous findings based on multiscale PDF analysis implicated the role of cascade dynamics in postural control to destabilizing task constraints [1]. We studied quiet standing in human participants holding a hollow tube filled with sand or water, with sand posing more stable load than water (Fig. 1). Quiet standing with the sand-filled tube evoked stronger cascade-like signatures of intermittency in medium log-timescales, but task constraints that destabilize quiet standing and demand compensatory adjustments increased non-Gaussianity at shorter log-timescales without any change in the decrease of non-Gaussianity at longer timescales. The result of this short-timescale destabilization is a non-Gaussianity curve with negative linear and cubic components along with a positive quadratic component—in contrast to the less-destabilizing sand-filled tube only prompting a negative quadratic non-Gaussianity curve (Fig. 2). Another way to phrase this point, especially to support subsequent remarks, is that the stabilization compensatory for novel destabilizing perturbations induced a ‘sign change’ in the polynomial form in how non-Gaussianity decays with log-timescale (i.e., negative linear and cubic terms alongside a positive quadratic).

**Fig. 1.**
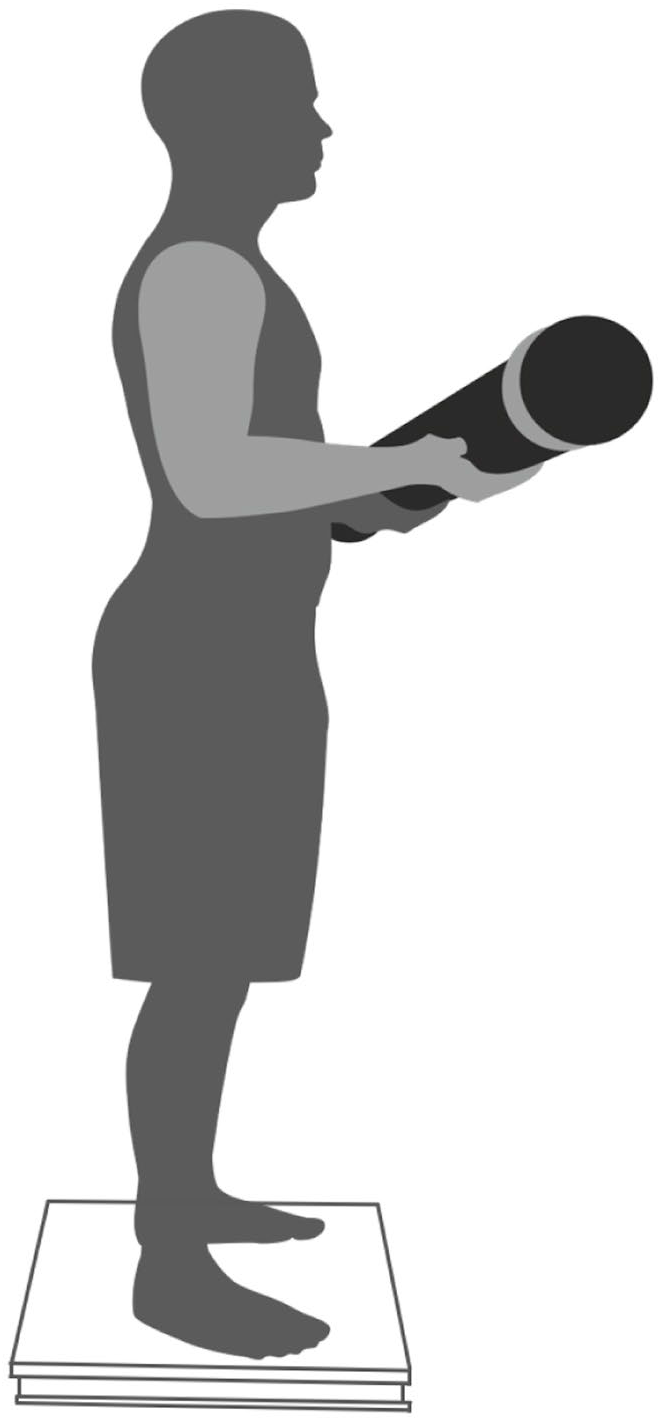
Each participant stood quietly while balancing a sand- or water-filled tube for 30 s.

**Fig. 2.**
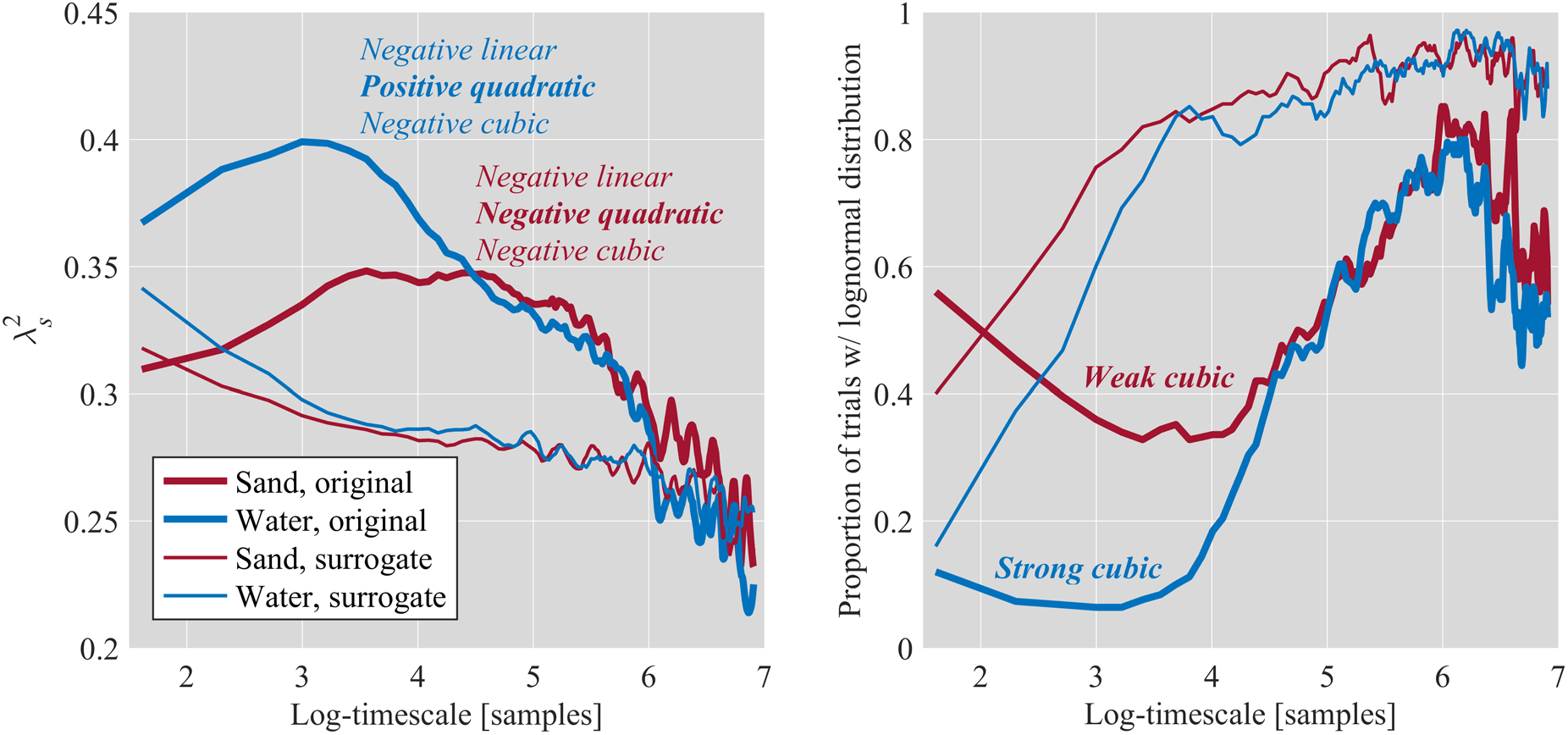
An illustration of log-timescale dependence of mean 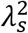 (left) and empirical proportion of trials with a lognormal distribution (right) for CoP planar displacement in participant standing quietly while balancing sand- and water-filled tubes for 30 s, as reported previously [1].

The present reanalysis’s overarching goal is to show that this polynomial form of the non-Gaussianity curve allows more subtle cascade dynamics modeling. Our prior work with these data only demonstrated the sign-changing effect of destabilizing constraints on posture. The present reanalysis aims to use this polynomial form to delve into this cascade’s structure to differentiate the sometimes monolithic-seeming ‘cascade dynamics’ into at least two segregable streams: one to absorb perturbation reactively and the other to respond proactively to maintain the task performance. Previous work examining non-Gaussianity using multiscale PDF analysis has shown that negative-linear and positive-quadratic components indicate scale-invariance cascades and scale-dependent cascades, respectively [29]. Scale-dependent task constraints may thus act primarily as a stimulus to strengthen the cascade-like dynamics at progressively shorter timescales and manifest in stronger positive quadratic (i.e., upwardly-opening parabolic) decay. On the other hand, human performance in executive control and attentional shifting of focus exhibits a widely scale-invariant form, similarly exhibiting branching, cascade-driven power-law forms from the finest grains of eye movements across a screen [30] to the medium term of remembering events both past [31] and planned [32,33], and to the longest timescales of organisms searching and reorganizing their environments [34]—even over multiple lifespans [35].

#### 1.3.1. Hypothesis-1: Reactive response to destabilizing task constraints would manifest as a positive-quadratic decay of non-Gaussianity with log-timescale for CoP

Our first hypothesis addresses how a scale-dependent cascade reflects a passive reaction to perturbing task constraints. In the present experimental setting of asking participants to stand quietly, the participant’s intentional postural control might introduce a relatively scale-invariant cascade to complement the scale-dependent cascade responding to the task constraints. The multiscale PDF analysis thus opens up the possibility of testing a postural control model in which the organism embodies a form of cascade dynamics that can engender, at least, both reactive scale-dependent cascades and proactive scale-invariant cascades. Specifically, we predict that a positive quadratic term would be specifically reactive to the more destabilizing constraints. In other words, standing with a water-filled tube would generate non-Gaussianity profiles showing a stronger positive quadratic relationship with log-timescale than standing with a sand-filled tube (Hypothesis-1). As noted, later on, this reactive scale-dependent cascade will be moderated by rambling sway, but this hypothesis is to establish that the cascade will appear in CoP in response specifically to the task constraint without explicitly intentional equilibrium-point control.

#### 1.3.2. Hypothesis-2: Proactive response to destabilizing task constraints would manifest as linear decay in non-Gaussianity with log-timescale specifically for rambling sway

Pinpointing a proactive scale-invariant cascading process that stabilizes posture requires a closer examination of postural control. Specifically, postural stabilization requires resetting the equilibrium points at which the organism can stand—the CoP might wobble randomly, but the CoP trajectory reflects both intentional and random postural shifts around the base of support. A compelling method to parse these differences from the CoP trajectory is the rambling/trembling decomposition [2,3]. In this framework, the rambling sway reflects cumulative displacement to new equilibrium points, whereas the trembling components reflect the deviations of posture from the new equilibrium points. Between rambling and trembling sway, it is the rambling sway that carries the systematic effects of age [36,37], injury and disease [38,39], and task [40,41]. Hence, we hypothesize that rambling sway would carry the cascade-dynamics proactively working to reset equilibrium, and so we predict that the negative linear component of non-Gaussianity decay across log-timescale should appear in rambling sway (Hypothesis-2). We expected no differences in the capacity of trembling non-Gaussianity to exhibit the linear or quadratic effects of log-timescale different from the CoP non-Gaussianity.

We can situate rambling and trembling in the same framework of intermittency that we had described CoP initially. We described intermittency conceptually as a system’s capacity to hover close to an attractive set of task-satisfying behaviors without getting locked into perfectly regular task satisfaction. Intermittent behaviors remain fluid enough to wander towards more extreme behaviors. Hence, if behavior follows a bell-shaped curve, cascade-driven intermittency is the fluid spreading of behavior from ‘task satisfaction’ in the bulky middle into ‘deviations from the task’ in the extreme tails. The rambling-trembling decomposition requires elaborating this metaphor to allow the bell-shaped curve itself to move. The rambling of posture allows task satisfaction to take on different equilibrium points. Hence, the CoP intermittency might reflect deviations around one quiet stance and deviations among and around multiple distinct stances. Our previous analysis of this data only used non-Gaussianity on the single distribution of CoP [1]. This reanalysis tests non-Gaussianity of two other different distributions: one describing the rambling across the base of support and the other describing the trembles above and beyond each new equilibrium point.

#### 1.3.3. Hypothesis-3: Multifractal nonlinearity supports the segregation of CoP fluctuations into divergent and asymmetric rambling vs. trembling sway

An immediate question for the previous portrayal pertains to whether the decomposition between rambling and trembling is consistent with cascade dynamics. After all, an important feature of cascade dynamics has been a reluctance to explain findings strictly in terms of reduced independent components [42–44]. Certainly, cascade dynamics offers the capacity to explain and predict intermittent behavior based on non-Gaussian, power-law relationships unfolding across many scales. The fractal dimensions governing these scaling relationships can be more robust and generic than independent components and constraints [45,46]. However, the fractal structure does not preclude the explanation and prediction of how behavior uses disparate functions. Cascade dynamics never prohibit diversity. On the contrary, diversity in cascading processes across timescales will engender a diverse set of structures supporting the observed behavior. This point is central to cascade dynamics: cascades are dramatically more likely to generate multifractal forms than forms with only one monolithic fractal structure.

The multifractal structure is a critical feature of cascade dynamics, allowing heterogeneous bodies to flexibly and adaptively perform heterogeneous behaviors. For instance, we had noted above that the executive control and attentional processes exhibit a fractal organization. Now, we note that variation in this fractal structure supports qualitative changes in executive functioning. That is to say, these executive processes reflect multifractal organization. The temporal variation in the fractal structure of human behavior supports our flexible use of qualitatively different rules in executive function tasks [47–50]. Human behavior also exhibits spatial variation in fractal structure, and subtle differences in fractal structure across the body mediate the pickup of ambient information by disparate body parts to serve different perceptual and cognitive functions [51–54]. Hence, multifractality is sooner the diversity-promoting rule than an exotic exception.

Multifractal nonlinearity can refine our understanding of postural control. The primary usefulness of multifractality is modeling how well a cascade process can segregate into disparate streams. In this sense, multifractality is the geometrical guarantee that cascade dynamics are fluid without being holistic, resisting reduction to anatomical parts without being monolithic. Of course, fluidity can seem like holism; resistance to reduction can seem like monolithic status. However, the critical point here is that multifractality can reflect nonlinear interactions across scales, and this nonlinearity can prompt divergence without requiring the model to add up fundamentally separate parts. We expect that multifractality would support the segregation of cascade-driven behavior into disparate streams. Hence, we predict that multifractal nonlinearity would effectively decouple the non-Gaussianity of CoP as a whole from the non-Gaussianity of rambling and trembling sway, in that non-Gaussianity of CoP would grow inversely with that of its rambling and trembling components (Hypothesis-3a).

Multifractality is classically a guarantee of asymmetry, not just across time but across space [55,56]. So, whereas our previous work only examined planar CoP displacements and did not distinguish between anterior-posterior (AP) and medial-lateral (ML) axes of sway, the rambling-trembling decomposition requires parsing CoP variability separately along the AP and ML axes. We assume that the rich internal structure composing CoP offers ample capacity for intermittency in both directions. Thus, we predict no differences in CoP non-Gaussianity between these axes. However, CoP decomposition produces rambling and trembling sway, which, as subsets of the measured CoP series, necessarily carry less internal structure. Consequently, we expect intermittency along one axis to come at the expense of intermittency along the other one, leading non-Gaussianity measures in AP rambling or trembling to vary inversely with those in ML rambling or trembling, respectively (Hypothesis-3b).

#### 1.3.4. Hypothesis-4: Multifractal nonlinearity moderates the segregation of postural control into scale-invariant and scale-dependent cascades specific to rambling sway

Multifractal nonlinearity moderates all influences of task constraints and intentional control on cascade dynamics. Indeed, extensive research has shown that multifractal nonlinearity of upright posture changes in response to task constraints [57] and intentions [58], and changes in multifractal nonlinearity influences participants’ perceptual judgments [53,59,60]. In the present case, we hypothesize that multifractal nonlinearity would shape all features articulated above: the scale-dependent cascade carrying the reactive response to destabilization, the scale-invariant cascade carrying proactive stabilization specifically under destabilizing task constraints, and the reconfiguration of cascade dynamics supporting intentional resetting of equilibrium points in general (Hypothesis-4).

The influence of multifractal nonlinearity on each of these features warrants specific predictions about the profile of non-Gaussianity across log-timescale. As for the scale-dependent cascade carrying the reactive response to destabilization, holding the water-filled tube strengthens the positive quadratic profile in non-Gaussianity with log-timescale [1]. We now predict that multifractal nonlinearity would support this positive quadratic component in rambling sway (Hypothesis-4a) accentuating the expected replication of effects on CoP (Hypothesis-1). Similarly, we had noted above that the water-filled tube strengthened the negative linear effect of log-timescale, indicating a scale-invariant cascade associated with proactive stabilization. We now predict that multifractal nonlinearity would support this negative linear effect specifically for rambling sway when holding the water-filled tube (Hypothesis-4b). The rambling-specificity of proactive response may initially seem more natural than that of reactive response. Indeed, the term ‘reactive’ might sound less deliberate and potentially more directionless and more relevant to trembling. However, the reaction to scale-dependent constraints is no less of an intentional engagement of movement degrees of freedom. For instance, the reactive catching of a ball thrown from afar is no less task-sensitive than throwing it to a distant location. Similarly, the reaction to postural task constraints is an essential factor in the task-sensitive decision to maintain or reset equilibrium points.

Finally, the foregoing allows us to take a new perspective on our previous findings and propose a more elaborate portrayal of cascade dynamics supporting postural control. First, as noted above, our previous work had documented that the multifractal nonlinearity supported the water-filled tube’s capacity to prompt stronger negative linear and cubic components as well as a significant positive quadratic components—a departure from the negative quadratic effect in the non-Gaussianity curve under the less destabilizing constraint of the sand-filled tube. In retrospect, although these results suggested that more stringent task constraints can prompt novel cascade dynamics, they gave no voice to any postural dynamics rising to the no-less-contrived task of holding a sand-filled tube. Secondly, that work had operated on CoP planar displacements. Planarity was not a significant constraint because previous work did not press any predictions regarding directionality. However, by committing to a non-directional investigation of CoP fluctuations, the previous work necessarily omitted the more theoretically weighty intentionality issue. Indeed, though more stable than a water-filled tube, a sand-filled tube remains a destabilizing task demand requiring resetting equilibrium points above and beyond an unencumbered quiet stance.

The rambling-trembling decomposition now offers us the capacity to test whether the sign-changed polynomial in non-Gaussianity is a generic feature to the intentional resetting of equilibrium points in postural control. That is, beyond merely predicting a piecewise addition of non-rambling quadratic and rambling linear components in the case of the water-filled tube (i.e., Hypotheses-4a and 4b), we predict that the sign-change of the non-Gaussianity polynomial (previously only attributed to the water-filled tube) is generic to both task manipulations (i.e., both fillings of the tube) but specifically moderated by rambling sway. Indeed, past results suggest that the general negative-linear and positive-quadratic effects on CoP non-Gaussianity profiles might be stronger while holding the water-filled tube— sufficiently strong to appear in our previous evidence of planar CoP displacements. However, now we predict that multifractal nonlinearity interacts with rambling sway to enact the sign-change in the polynomial for postural control while holding the sand-filled tube as well (i.e., significantly negative, positive, and negative coefficients for the linear, quadratic, and cubic polynomials of log-timescale, respectively, for original series’ rambling components independent of the task constraints; Hypothesis-4c). Finally, we predict that multifractal nonlinearity would moderate this sign-changed polynomial in the original series’ rambling components (Hypothesis-4d).

Our emphasis in the present work is to suggest that rambling sway is unique in carrying the reactive and proactive components of cascade dynamics. Therefore, we do not predict that these multifractal effects on CoP or trembling would support the sign change needed to articulate negative-linear and positive-quadratic terms indicating the segregation of scale-invariant vs. scale-dependent cascades.

#### 1.3.5. Hypothesis-5: The ambiguity of lognormal-like tail heaviness across scales in the case of rambling sway

To situate these abstract distinctions in more concrete terms, the preceding four hypotheses all entail non-Gaussianity of rambling sway, but they amount to an expectation of heavy tails only some of the time. For instance, the expectations of negative linear and positive quadratic terms amount to strong non-Gaussianity at small scales—but not of heavy tails. On the contrary, whereas trembling proceeds throughout CoP sway, rambling is, by definition, a mix of sustained sway punctuating mostly sustained lack of sway. So, for short timescales, rambling sway is sparser, and so the resulting bell-shaped PDF curve is more likely to have a taller, wider midsection decaying more steeply towards thinner tails. Hence, lognormality would be too scarce for short timescales, and the larger displacements in rambling between equilibrium points will only contribute to a heavy-tailed lognormal distribution at longer timescales long enough to capture these more sustained movements.

We predict that the abrupt transitions in rambling sway (i.e., alternating between near-zero magnitude to large equilibrium-point resettings) would produce an equally abrupt transition in the proportion of lognormal rambling displacement PDFs across timescales. Specifically, we predict that rambling sway would show a positive cubic relationship of lognormality with log-timescale (Hypothesis-5). Indeed, we previously found a cubic form for planar CoP displacements’ lognormality relationship with log-timescale, with the more destabilizing task constraint (i.e., the water-filled tube) yielding a stronger cubic relationship [1]. Consistent with the rest of our present reanalysis, we aim here to anchor effects of the intentional response to task constraints in rambling sway. Therefore, we expect that rambling sway would show a stronger positive cubic relationship in general than CoP or trembling sway because rambling sway would exhibit very thin tails for small log-timescale and heavier lognormal-like tails only for much larger log-timescale (Hypothesis-5a). For the same reason, we also expect that rambling sway would show stronger cubic relationships for the water-filled tube (Hypothesis-5b). Because this tail heaviness of lognormality emphasizes extreme events and not the internal cascade-driven structure of sway, we expect lognormality along the AP and ML axes to vary inversely as extreme events in one direction come at the expense of the other (Hypothesis-5c).

## 2. Methods

### 2.1. Participants

Ten adult men (*mean±1SD* age = 21.4±1.1 years) with no neurological, muscular, or orthopedic conditions participated in this study after providing institutionally-approved informed consent.

### 2.2. Experimental setup and procedure

The participants stood quietly on a force platform (AMTI Inc., Watertown, MA) with their feet shoulder-width apart and bimanually balanced tubes half-filled with sand or water (*l×d* = 150×20 cm, *m* = 8 kg; Fig. 1). The participants were instructed to balance the tubes with as much stability as possible. Balancing the water-filled tube thus imposed more stringent postural constraints than balancing the sand-filled tube. The ground reaction forces were recorded for 30-s trials at 100 Hz. Each participant completed five trials per task, with trial order pseudo-randomized across participants.

### 2.3. Rambling and trembling decomposition

Trial-by-trial ground reaction forces yielded a 2D foot center of pressure (CoP) series, each dimension describing CoP position along anterior-posterior (*AP*) and medial-lateral (*ML*) axes (CoP_*AP*_ and CoP_*ML*_, respectively). The rambling and trembling components of CoP along the *AP* and *ML* medial-axes are computed based on Zatsiorsky and Duarte’s method [2]. This method begins by identifying instant equilibrium points (IEPs) as instances when the horizontal ground reaction force along the *AP* and *ML* axes (Force_*AP*_ and Force_*ML*_, respectively) changes its sign. CoP at all IEPs are determined and interpolated by local linear interpolation of the Force_*AP*_ time-history data using the cubic spline function to obtain the rambling trajectories along the *AP* and *ML* axes (Rambling_*AP*_ and Rambling_*ML*_, respectively). When using cubic splines, each segment between the IEPs is connected by a 3^rd^-order polynomial, and the slope of each cubic polynomial is matched at these points [61]. CoP deviation from the respective rambling trajectory is computed to obtain the trembling trajectories along the *AP* and *ML* axes (Trembling_*AP*_ and Trembling_*ML*_, respectively). The first derivative of the obtained CoP_*AP*/*ML*_, Rambling_*AP*/*ML*_, and Trembling_*AP/ML*_ series served as displacement or fluctuation series.

### 2.4. Assessing multifractality due to multiplicativity

#### 2.4.1. Direct estimation of multifractal spectrums

Chhabra and Jensen’s direct method [62] was used to estimate multifractal spectrum widths *Δα* of each original CoP_*AP/ML*_, Rambling_*AP/ML*_, and Trembling_*AP/ML*_ displacement series. This method samples series *u*(*t*) at progressively larger scales such that the proportion of signal *P_i_*(*L*) falling within the *i^th^* bin of scale *L* is

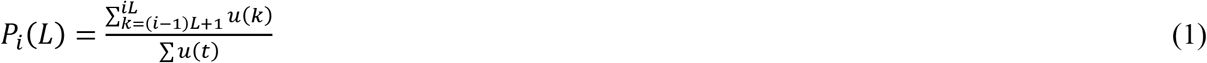

As *L* increases, *P_i_*(*L*) represents progressively larger proportion of *u*(*t*),

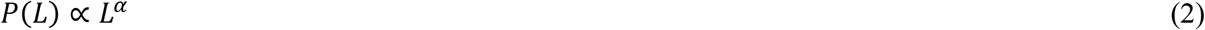

suggesting growth of proportion according to one “singularity” strength *α* [56]. *P*(*L*) exhibits multifractal dynamics when it grows heterogeneously across time scales *L* according to multiple singularity strengths, such that

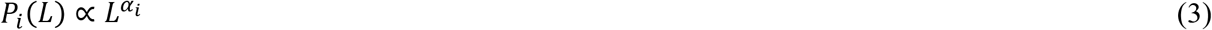

whereby each *i^th^* bin may show a distinct relationship of *P*(*L*) with *L*. The width of this singularity spectrum, *Δα*(*α_max_* – *α_min_*), indicates the heterogeneity of these relationships [24,63].

Chhabra and Jensen’s method [62] estimates *P*(*L*) for *N_L_* nonoverlapping bins of *L*-sizes and transforms them into a “mass” *μ* using a *q* parameter emphasizing higher or lower *P*(*L*) for *q* >1 and *q* < 1, respectively, as follows

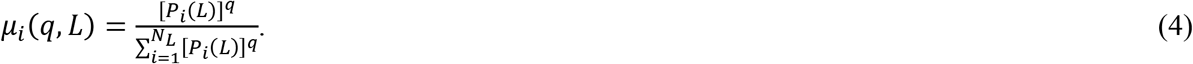

*α*(*q*) is the singularity for mass μ(q)-weighted P(L) estimated by

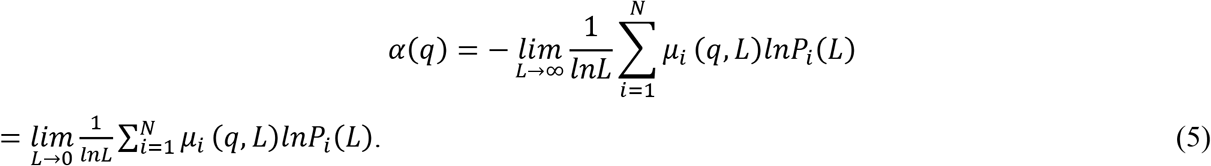

Each estimated value of *α*(*q*) belongs to the multifractal spectrum only when the Shannon entropy of *μ*(*q, l*) scales with *L* according to the Hausdorff dimension *f*(*q*), where

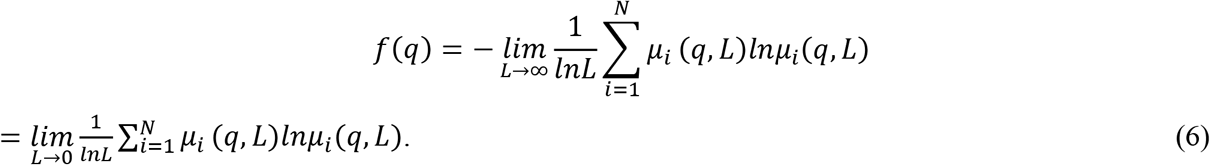

For values of *q* yielding a strong relationship between Eqs. (5 & 6)—in this study, exhibited a correlation coefficient, *r* > 0.9, the parametric curve (*α*(*q*), *f*(*q*)) or (*α, f* (*α*)) constitutes the multifractal spectrum.

#### 2.4.2. Surrogate testing

To identify whether nonzero *Δα* reflected multifractality due to nonlinear interactions across timescales, *Δα* of each original series was compared to *Δα* of the corresponding Iterated Amplitude Adjusted Fourier Transformation (IAAFT) surrogates [64,65]. IAAFT randomizes original values time-symmetrically around the autoregressive structure, generating surrogates that randomize phase ordering of the series’ spectral amplitudes while preserving linear temporal correlations. If *Δα* for the original series exceeds a 95% confidence interval (CI) of *Δα* for 32 IAAFT series (i.e., *P* < 0.05), then the original series shows multifractality due to multiplicativity, which is quantifiable using the one-sample *t*-statistic (henceforth, *t*_MF_) comparing *Δα* for the original series to that for the 32 surrogates.

### 2.5. Multiscale probability density function (PDF) analysis

We subjected each original CoP_*AP/ML*_, Rambling_*AP/ML*_, and Trembling_*AP/ML*_ displacement series and corresponding IAAFT surrogate to multiscale PDF analysis. Multiscale PDF analysis characterizes the distribution of abrupt changes in CoP PED series {*b*(*t*)} using the PDF tail. The first step is to generate {*B*(*t*)} by integrating {*b*(*t*)} after centering by the mean *b_ave_* (Fig. 3, top):

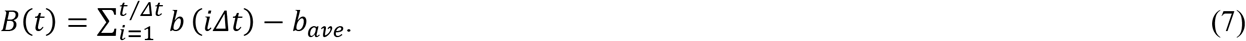

**Fig. 3.**
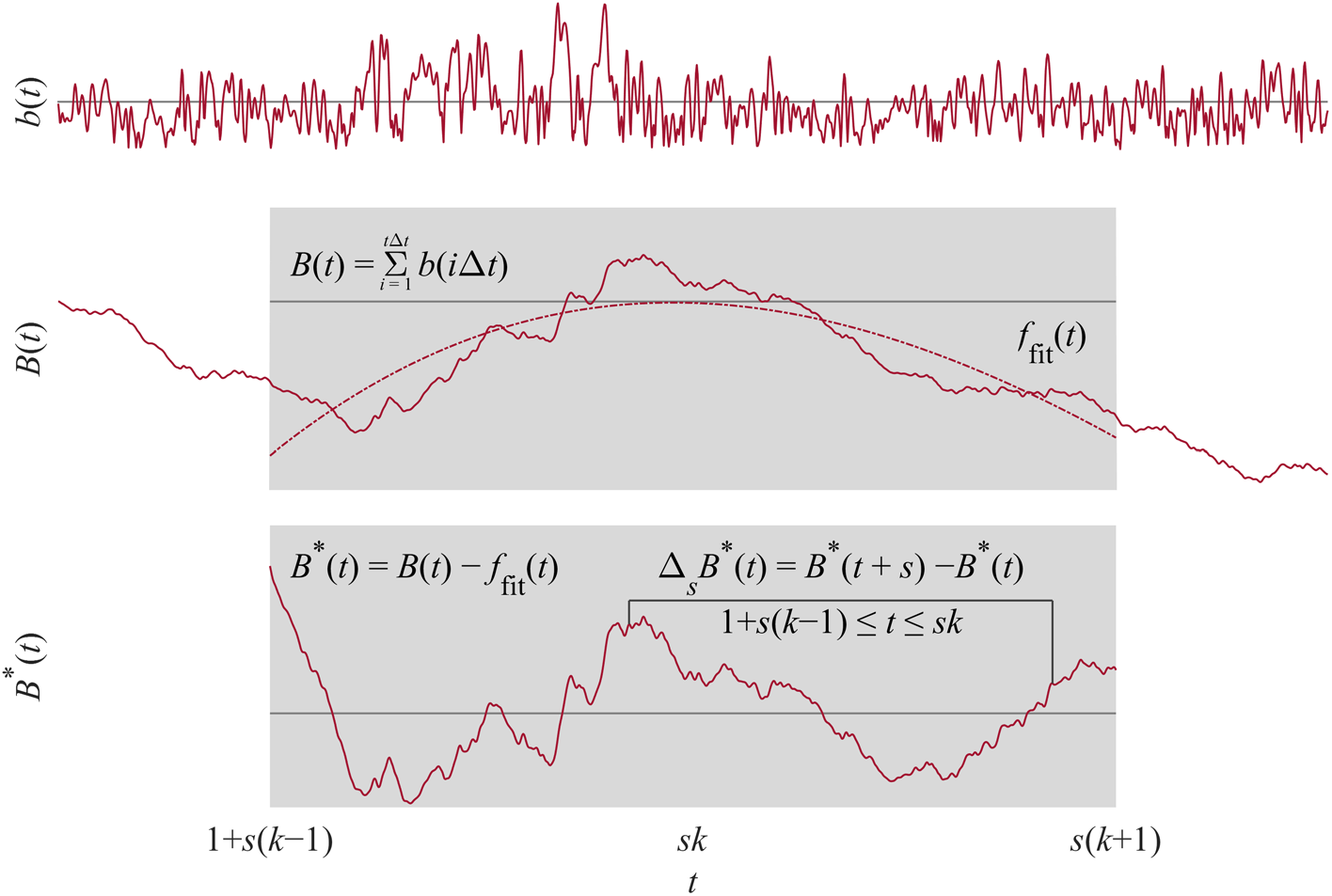
Schematic illustration of multiscale probability density function (PDF) analysis. Top: The first step is to generate {*B*(*t*)} by integrating {*b*(*t*)} after centering by the mean 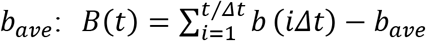. Middle: A a 3^rd^ order polynomial detrends {*B*(*t*)} within *k* overlapping windows of length 2*s, s* being the timescale. Bottom: Intermittent deviation *Δ_s_B*(*t*) in *k^th^* window from 1 + *s*(*k* – 1) to *sk* in the detrended time series {*B^d^*(*t*) = *B*(*t*) – *f_fit_*(*t*)} is computed as *Δ_s_B^d^*(*t*) = *B^d^*(*t* + *s*) – *B^d^*(*t*), where 1 + *s*(*k* – 1) ≤ *t* ≤ *sk* and *f_fit_*(*t*) is the polynomial representing the local trend of {*B*(*t*)}, of which the elimination assures the zero-mean probability density function in the next step.

A a 3^rd^ order polynomial detrends {*B*(*t*)} within *k* overlapping windows of length 2*s, s* being the timescale (Fig. 3, middle). Intermittent deviation *Δ_s_B*(*t*) in *k^th^* window from 1 + *s*(*k* – 1) to *sk* in the detrended time series {*B^d^*(*t*) = *B*(*t*) – (*t*)} is computed as *Δ_s_B^d^*(*t*) = *B^d^*(*t* + *s*) – *B^d^*(*t*), where 1 + *s*(*k* – 1) ≤ *t* ≤ *sk* and *f_fit_*(*t*) is the polynomial representing the local trend of {*B*(*t*)}, of which the elimination assures the zero-mean probability density function in the next step (Fig. 3, bottom). Finally, *Δ_S_B* is normalized by the SD (i.e., variance is set to one) to quantify the PDF.

To quantify the non-Gaussianity of *Δ_S_B* at timescale *s*, the standardized PDF constructed from all the *Δ_s_B*(*t*) values is approximated by the Castaing model [66], with *λ_s_* as a single parameter characterizing the non-Gaussianity of the PDF. *λ_s_* is estimated as

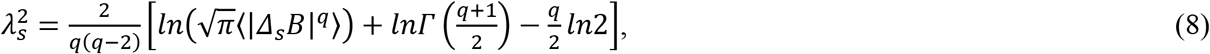

where 〈|*Δ_s_B*|^*q*^〉 denotes an estimated value of *q^th^* order absolute moment of {*Δ_S_B*}. As 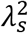 increases, the PDF becomes increasingly peaked and fat-tailed (Fig. 4, left). 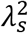 can be estimated by Eq. (8) based on *q^th^* order absolute moment of a time series independent of *q*. Estimating 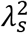 based on 0.2^th^ moment (*q*= 0.2) emphasizes the center part of the PDF, reducing the effects of extreme deviations due to heavy-tails and kurtosis. We used 0.2^th^ moment because estimates of 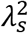 for a time series of ~ 3000 samples are more accurate at lower values of *q* [21].

**Fig. 4.**
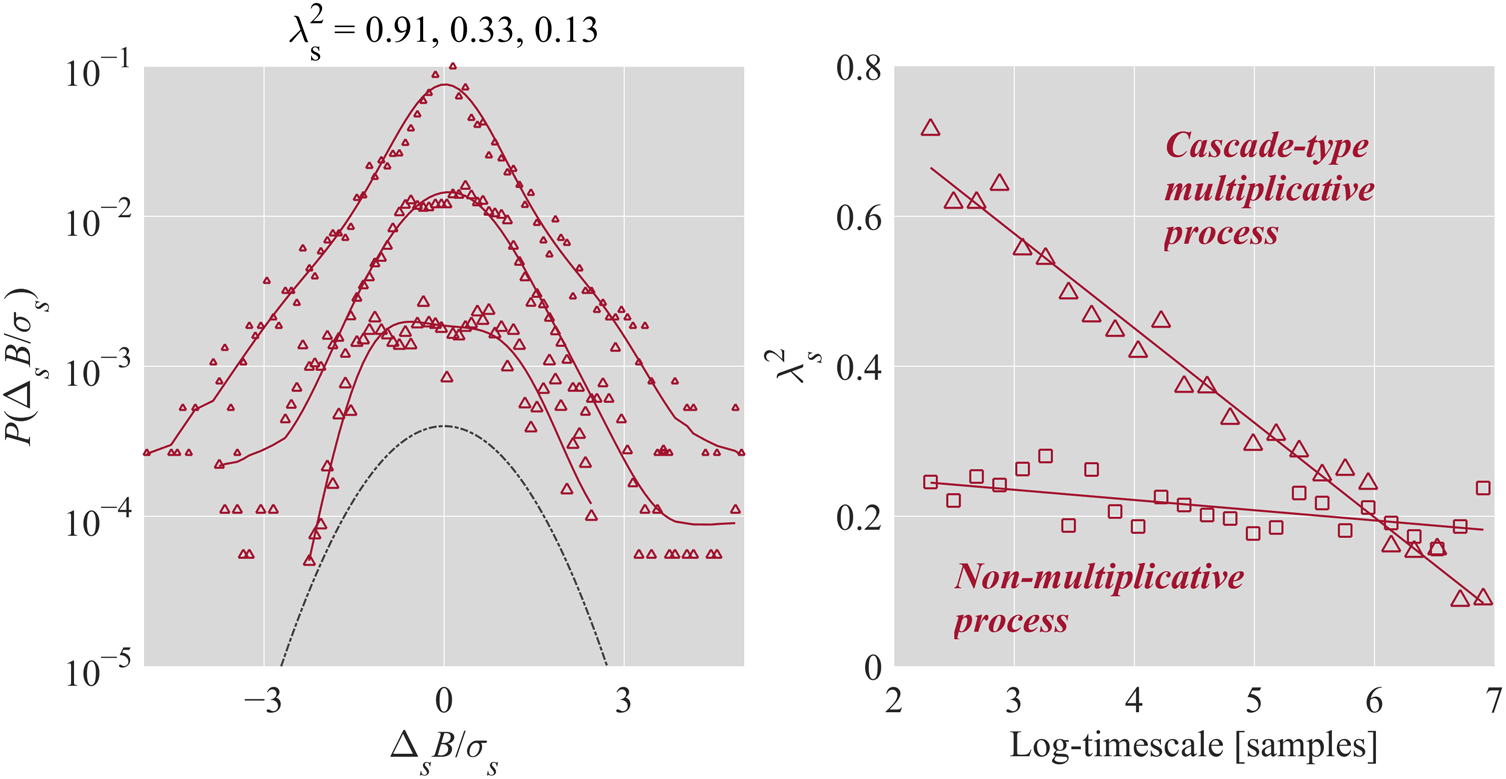
Schematic illustration of non-Gaussianity index *λ_s_*. Left: The relationship between 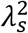 and shapes of PDF plotted in linear-log coordinates for 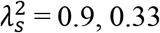, and 0.13 (from top to bottom). For clarity, the PDFs have been shifted vertically. As 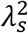 increases, the PDF becomes increasingly peaked and fat-tailed. As *λ_s_* decreases, the PDF increasingly resembles the Gaussian (dashed line), assuming a perfect Gaussian as 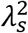 approaches 0. Right: Cascade-type multiplicative processes yield the inverse relationship 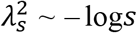.

Cascade-type multiplicative processes yield the inverse relationship 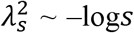 (Fig. 4, right) [20]. For the present purposes, we quantified 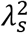 for each original CoP_*AP/ML*_, Rambling_*AP/ML*_, and Trembling_*AP/ML*_ displacement series and corresponding IAAFT surrogate at timescales of 5 to 1000 samples (i.e., 50 ms to 10 s) at steps of 5 samples (50 ms). Using IAAFT surrogate here allowed identifying non-Gaussianity attributable to linear structure (i.e., the histogram and linear autocorrelation of displacement series). Departures of the original series’ non-Gaussianity from that for corresponding surrogates indicate the degree to which non-Gaussianity reflected the nonlinear interactions across scales characteristic of cascade dynamics.

Multiscale PDF analysis of non-Gaussianity and traditional MLE methodology using Akaike Information Criterion (AIC) differ in their emphasis. In contrast to traditional MLE, which is disproportionately sensitive to the less-populated tails, multiscale PDF analysis addresses the central, better-populated bulk of the PDF. Hence, we also included MLE to complement multiscale PDF analysis, specifically to portray the analytical effect of sensitivity to different portions of the distribution. We used AIC to test which among a power law, lognormal, exponential, or gamma distribution characterized the PDF.

### 2.6. Statistical analysis

To test for crossovers in non-Gaussianity, a linear mixed-effects (LME) model using *lmer* function [67] in R package *lme4* tested 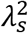 vs. log-timescale curves for orthogonal linear, quadratic, and cubic polynomials, for their interactions with grouping variables (Axis × Variable × Task × Original, where Variable encoded Rambling and Trembling as compared to CoP and where Original encoded differences in original series from surrogates) and with *t*_MF_. Statistical significance was assumed at the alpha level of 0.05 using R package *lmerTest* [68]. To test how lognormality changed with log-timescale, a generalized linear mixed-effects (GLME) model fit changes in Lognormality as a dichotomous variable (Lognormal = 1 vs. Exponential = 0) using orthogonal linear, quadratic, and cubic polynomials and tested interaction effects of grouping variables (Axis × Variable × Task × Original) with those polynomials using glmer [69] in *lme4*.

## 3. Results

Figs. 5 exemplifies the relationships between the horizontal ground reaction force and CoP along the *AP* and *ML* axes, and its rambling and trembling components, in a representative participant standing quietly while balancing sand- and water-filled tubes for 30 s. Fig. 6 provides a schematic of multiscale PDF characterization of postural sway in representative participants balancing sand- and water-filled tubes for 30 s.

**Fig. 5.**
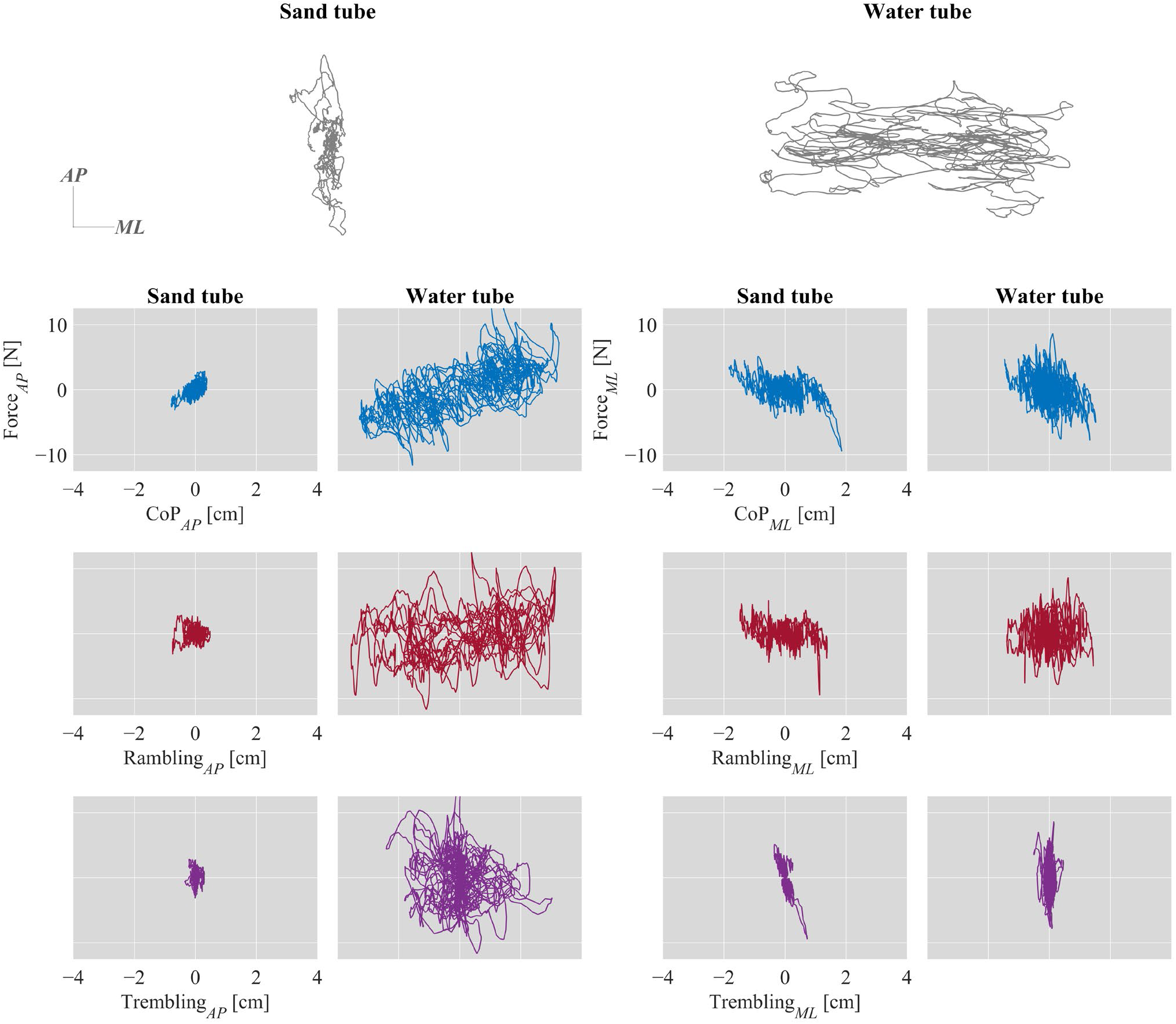
Relationships between the horizontal ground reaction force and CoP along the *AP* and *ML* axes, and its rambling and trembling components, in a representative participant standing quietly while balancing sand- and water-filled tubes for 30 s.

**Fig. 6.**
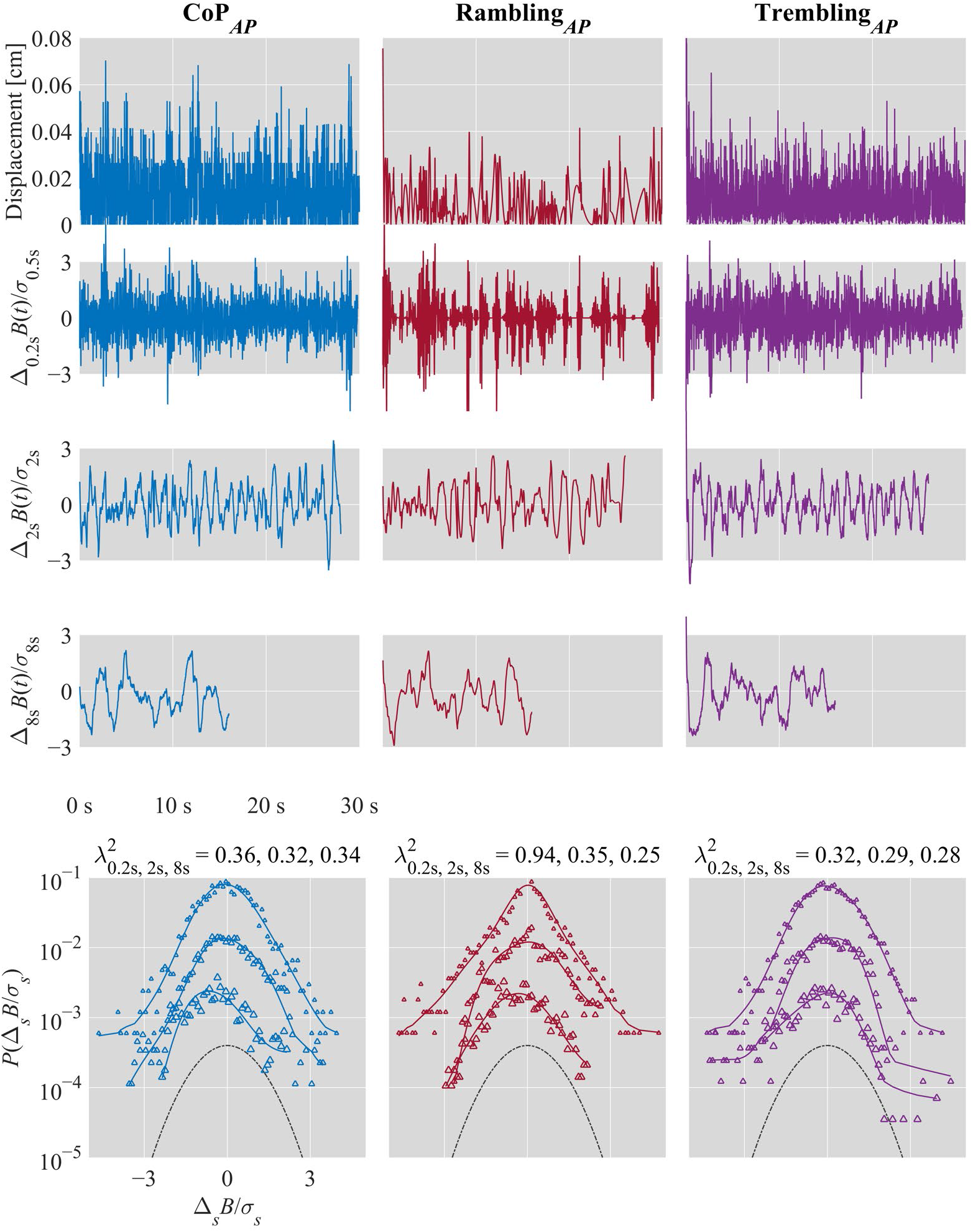
Multiscale PDF characterization of postural sway in a representative participant standing quietly while balancing sand-filled tube for 30 s. From left to right: CoP, rambling, and trembling trajectories along the *AP* axis. From top to bottom: Displacement series. {*Δ_s_B*(*i*)} for *s* = 0.2, 2, and 8 s; Standardized PDFs (in logarithmic scale) of {*Δ_s_B*(*i*)} for *s* = 0.2, 2, and 8 s (from top to bottom), where *σ_s_* denotes the *SD* of {*Δ_s_B*(*i*)}.

### 3.1. Testing Hypothesis-1: Positive quadratic decay in non-Gaussianity with log-timescale in reactive response to destabilizing task constraints

Original CoP series showed a significantly stronger positive-quadratic decay in non-Gaussianity with log-timescale in response to the water-filled tube (*B* = 2.56×10^0^, *P* = 0.044; Fig. 7, top left).

**Fig. 7.**
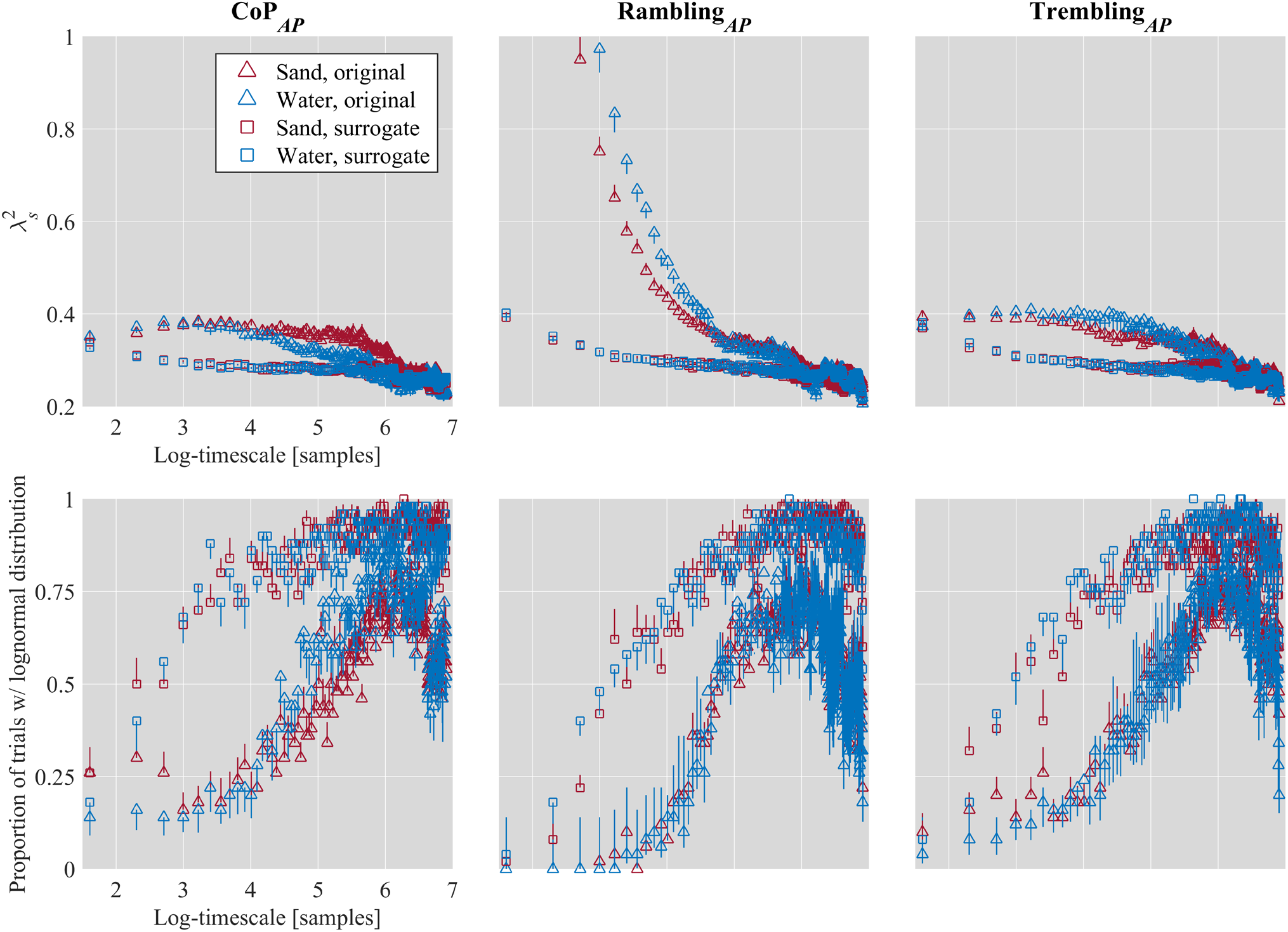
Log-timescale dependence of mean 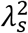 (top; Sections 3.1 to 3.4; Table S1) and empirical proportion of trials with a lognormal distribution (bottom; Section 3.5; Table S2), for the CoP, rambling, and trembling trajectories along the *AP* axis in the postural tasks of standing quietly while balancing sand- and water-filled tubes for 30 s. Vertical bars indicate ±1*SEM* (*N* = 10).

### 3.2. Testing Hypothesis-2: Negative linear decay in non-Gaussianity for rambling sway in proactive response to destabilizing task constraints

The original rambling series in the *AP* direction showed a significantly negative-linear decay in non-Gaussianity with log-timescale in response to the water-filled tube (*B* = –10.77×10^0^, *P* < 0.001). The original rambling series in the *ML* direction exhibited a reversal of this linear component (*B* = 1.89×10^1^, *P* < 0.001; Fig. 7, top center).

### 3.3. Testing Hypothesis-3: Multifractal nonlinearity moderated the segregation of CoP sway into rambling and trembling sway

#### 3.3.1. Testing Hypothesis-3a: Multifractal nonlinearity supported an inverse relationship between non-Gaussianity in CoP and non-Gaussianity of rambling and trembling components of sway

Multifractal nonlinearity moderated significant deviations of rambling and trembling sway from CoP sway in the intercepts of non-Gaussianity profiles with log-timescale. In response to the sand-filled tube, multifractal nonlinearity led the original CoP series to exhibit significantly more non-Gaussianity than the surrogates along the *AP* axis (*B* = 4.60×10^−5^, *P* < 0.001; Fig. 7, top left). However, multifractal nonlinearity led rambling (*B* = –3.98×10^−5^, *P* < 0.001; Fig. 7, top center) and trembling (*B* = –3.55×10^− 5^, *P* < 0.001; Fig. 7, top left) sway both to show less non-Gaussianity along the *AP* axis. The water-filled tube reversed this pattern of effects, yielding CoP sway interacting with multifractal nonlinearity to produce less non-Gaussianity along the *AP* axis (*B* = –7.56×10^−5^, *P* < 0.001) but yielding rambling (B = 1.14×10^−4^, P < 0.001) and trembling (*B* = 1.13 10 ^4^. *P* < 0.001) sway with more non-Gaussianity.

#### 3.3.2. Testing Hypothesis-3b: Multifractal nonlinearity supported asymmetry in rambling and trembling sway

Multifractal nonlinearity led non-Gaussianity of CoP sway to increase even more so along the *ML* axis (*B* = 5.23×10^−5^, *P* < 0.001) with no difference due to the water-filled tube. However, non-Gaussianity in rambling and trembling sway showed that multifractal nonlinearity moderated opposite changes along the *ML* and *AP* axes. With greater multifractal nonlinearity, trembling sway was more non-Gaussian for the sand-filled tube (*B* = 2.03×10^−4^, *P* < 0.001), though rambling showed no difference from CoP in this case. The water-filled tube conspired with multifractal nonlinearity to produce significantly weaker non-Gaussianity in both rambling (*B* = 1.02×10^−4^, *P* < 0.001) and trembling (*B* = – 1.64×10^−4^, *P* < 0.001) sway along the *ML* axis than along the *AP* axis.

### 3.4. Testing Hypothesis-4: Multifractality nonlinearity moderated the segregation of cascade dynamics across rambling and trembling sway

#### 3.4.1. Testing Hypothesis-4a: Multifractal nonlinearity moderated the positive-quadratic decay in non-Gaussianity under destabilizing task constraints

In quiet standing with the water-filled tube, multifractal nonlinearity supported a significantly stronger positive quadratic effect of log-timescale for CoP sway (*B* = 1.15×10^−2^, *P* = 0.005). There were no differences in effects of multifractal nonlinearity on the quadratic log-timescale term for rambling or trembling sway specific to the water-filled tube. That is to say, multifractality contributed to the expression of scale-dependent cascades in posture even independently of explicitly intentional equilibrium-point control.

#### 3.4.2. Testing Hypothesis-4b: Multifractal nonlinearity moderated the negative-linear decay in non-Gaussianity under destabilizing task constraints specifically for rambling sway

In quiet standing with the water-filled tube, multifractal nonlinearity supported a significantly stronger negative linear effect of log-timescale for rambling sway (*B* = 1.30 10 ^2^, *P* = 0.011). This effect accentuated a lower-order significant negative interaction of multifractal nonlinearity with the linear term for the sand-filled tube (*B* = 3.63 10 ^2^, *P* < 0.001).

#### 3.4.3. Testing Hypothesis-4c: Multifractal nonlinearity moderated the polynomial components of non-Gaussianity vs. log-timescale relationship by inducing a sign change across all three components of the cubic polynomial for rambling sway

The original rambling series for the sand-filled tube showed the polynomial sign change of non-Gaussianity with log-timescale, that is, negative linear (*B* = –1.49×10^1^, *P* < 0.001), positive quadratic (*B* = 2.64×10^1^, *P* < 0.001), and negative cubic (*B* = –1.71×10^1^, *P* < 0.001) components along the *AP* axis. There was no further change in these components for the water-filled tube. Instead, multifractal nonlinearity accentuated all components of the polynomial sign change for the sand-filled tube (*B*s = – 3.63×10^−2^, 5.96×10^−2^, and –2.37×10^−2^ for the linear, quadratic, and cubic components, respectively; all *P* < 0.001), with no further difference in these effects with the water-filled tube.

Polynomial profiles of non-Gaussianity in rambling sway showed a stronger sign change along the *ML* than *AP* axis for the quadratic (*B* = 1.09×10^1^, *P* < 0.001) and cubic (*B* = –5.72×10^0^, *P* = 0.006) components. Multifractality nonlinearity weakened the negative linear decay of non-Gaussianity in rambling sway along the *ML* axis (*B* = 3.10×10^−2^, *P* < 0.001) but showed no effects on the quadratic and cubic components.

#### 3.4.4. Testing Hypothesis-4d: Multifractal nonlinearity moderated the polynomial components of non-Gaussianity vs. log-timescale relationship without inducing a sign change between linear and quadratic components for CoP and trembling sway

All other significant effects on the non-Gaussianity of original COP series showed similar signs across the linear, quadratic, and cubic terms, ensuring that, as Figure 6 shows, trembling sway resembled CoP in having similar negative linear, quadratic, and cubic polynomial relationship with log-timescale, along both the *AP* and *ML* axes (Supplementary Table S1). Multifractal nonlinearity significantly moderated all polynomial contributions to the non-Gaussianity vs. log-timescale relationship for CoP and trembling sway, with rare exceptions along both the *AP* and *ML* axes.

Differences in these multifractal moderation of polynomial terms with the water-filled tube were sparse, appearing only along the *ML* direction (Supplementary Table S1).

### 3.5. Testing Hypothesis-5: Lognormal-like heavy tails are ambiguous evidence of task-dependent intermittency in postural control

#### 3.5.1 Testing Hypothesis-5a: Rambling showed less lognormality at shorter timescales and cubic growth of lognormality at longer timescales

Rambling sway contributed to significantly more positive cubic growth of lognormality with log-timescale (*B* = 109.72, *P* = 0.006), indicating much lighter tails in rambling PDFs (Fig. 7, bottom center) than in CoP PDFs (Fig. 7, bottom left). Rambling sway’s tail heaviness ramped up quickly for larger values of log-timescale (e.g., log-timescales > 4), generating a peak with greater convexity indicated by the negative quadratic effect of log-timescale on non-Gaussianity (*B* = –262.07, *P* < 0.001).

#### 3.5.2 Testing Hypothesis-5b: Water-filled tube accentuated rambling sway’s scale-dependent growth of lognormality

The water-filled tube strengthened the dependence of rambling PDF’s heavy tail on log-timescale, yielding significantly more positive cubic form (*B* = 184.48, *P* = 0.004) leading up to a convex peak with significantly more negative quadratic form (*B* = –204.63, *P* = 0.047; Fig. 7, bottom center). The water-filled tube further accentuated scale-dependent growth of lognormality on the linear effect of logtimescale. That is, lognormality was even more rare for the smallest log-timescale (*B* = –0.57, *P* < 0.001) and grew also according to a positive linear relationship (*B* = 209.40, *P* = 0.014).

#### 3.5.3 Testing Hypothesis-5c: Rambling PDF’s tail heaviness showed tradeoff between AP and ML sway

Tradeoffs in rambling PDF’s tail heaviness between *AP* and *ML* axes only appeared in the case of the water-filled tube. For example, positive quadratic (*B* = 129.95, *P* < 0.001) and cubic (*B* = 95.42, *P* = 0.006) terms appeared in the *ML* scale-dependence of lognormality with the water-filled tube, reversing the same effects on the *AP* direction of CoP with the water-filled tube (positive quadratic: *B* = –95.13, P < 0.001; positive cubic: *B* = –60.88, *P* = 0.013). Similarly, rambling with the water-filled tube along the *ML* axis canceled the previously noted accentuation of linear (*B* = –299.93, *P* = 0.004)and cubic (*B* = –266.41, *P* = 0.001) terms observed along the *AP* axis. Trembling along the *ML* axis showed a negative quadratic effect (*B* = –161.20, *P* = 0.002) canceling the positive quadratic effect along the *AP* axis (*B* = 105.18, *P* = 0.004). However, not all *ML* effects were clear reversals of *AP* effects. For example, trembling along the *ML* axis showed a significant negative cubic relationship (*B* = –205.89, *P* < 0.001) despite trembling along the *AP* axis showing a nonsignificant cubic effect. In short, there were no tradeoffs in trembling PDF’s tail heaviness between the *AP* and *ML* axes (Supplementary Table S2).

## 4. Discussion

The present work tested five hypotheses regarding the relationship of intermittency to the dexterity of postural control. In particular, these five hypotheses aim to use a measure of non-Gaussianity to model the cascade dynamics supporting the rambling sway responsible for resetting postural equilibrium points. Hypothesis-1 was that non-Gaussianity would reveal a specific quadratic profile moderating the effect of task constraints indicating a scale-dependent cascade in CoP sway. Hypothesis-2 was that non-Gaussianity would reveal a specific linear profile moderating the effect of proactive stabilization indicating a scale-invariant cascade in rambling sway. Hypothesis-3 was that multifractal evidence of nonlinear interactions across scales would moderate the segregation of CoP into rambling and trembling. Hypothesis-4 was that multifractal evidence of nonlinear interactions across timescales would moderate the segregation of cascade dynamics of rambling sway into scale-dependent and scale-invariant cascades. Hypothesis-5 was that, unlike non-Gaussian evidence for the foregoing hypotheses, lognormal-like heavy tails would provide only ambiguous evidence of intermittency, appearing only for large log-timescale.

The results supported all hypotheses, and we summarize these before outlining implications. First, the quadratic term’s significant interactions with water-filled tube provided evidence that destabilizing task constraints prompted a scale-dependent cascade in posture—separate from but not exclusive of intentional equilibrium-point control. Second, the significant interactions of linear and quadratic terms with rambling provided evidence that the decisions to maintain or shift postural equilibrium points depended on two types of cascades: scale-invariant cascades typical of executive control and scaledependent cascades in response to perturbation. Third, the nonlinear degree of multifractal correlations beyond linearity was an essential moderator of two nested differentiation in postural control: first, rambling vs. trembling and then, within rambling, scale-dependent vs. scale-invariant cascades. Lastly, supporting the notion that multi-scale PDF analysis offers a clear analytical advantage, the non-Gaussianity evidence was strongest precisely where the evidence of heavy tails was the weakest. These results have two major sets of implications. The first set of implications pertain to the task-specificity of intermittency in postural control. The second set of implications pertain to the modeling of cascade dynamics to explain dexterous goal-directed behavior.

### 4.1. Task specificity of intermittency in postural control

The present evidence of non-Gaussianity provides yet stronger confirmation that intermittency is not merely incidental or endogenous to postural sway but rather specific to task constraints. Fractal and multifractal signatures of intermittency have long interested physiological research. One consequence of this interest has been a concerted effort to locate the precise specific anatomical structures or local physiological processes responsible for generating these fractal fluctuations, endogenously and even in the so-called absence of stimulation or behavioral prompting [70–76]. Indeed, if we appreciate that (multi)fractal signatures of intermittency have any usefulness, it might make an anatomical sort of sense to ask which part is responsible for it. However, the very premise that we should seek an independent component locally responsible for producing the observable behavior is at odds with the theoretical development of the contrary premise that (multi)fractal signatures of intermittency reflect more a global organization. The recognition of a global organization does not necessarily threaten interest in any specific local effects [77,78]. The present results show that non-Gaussian signatures of intermittency are, in fact, task-specific and indicative of the interactions of events at multiple timescales, local to global. Any so-called ‘absences’ of stimulation or behaviors supporting intermittency might only be absent contingent on a given timescale. But we see the intermittency as sooner specific to the multi-scaled contact of an organism with its task setting than specific to any single set of tissues.

These results speak to a functional specificity of intermittency. Resistance to finding intermittency in a single set of tissues might sound initially like lazy holism and so poor science [79], but that potential criticism can only come from a perspective that presumes that independent causal mechanisms require independent components. The present work finds the task-specificity of intermittency, specifically in those ‘rambling’ components of sway known to support intentional resetting of postural equilibrium points. Indeed, this elaboration is important because previous work made this intermittency seem to depend too much on a specific kind of task constraint [1]. If intermittency was only responsive to the more destabilizing water-filled tube, then it would leave the appearance that intermittency was dependent on stimulus—then perhaps the sand-filled tube would really appear like the absence of some crucial stimulation. The scale-dependent and scale-invariant cascades that rambling sway exhibits now show us that it is not the presence or absence of a specific stimulus but rather the intentional coordination of an organism with the task settings that generate intermittency. Hence, we avoid anchoring intermittency in the local anatomical part or the stimulation and, instead, anchor it in the branching, multiscaled relationships between the goal-directed organism and its environment.

### 4.2. Specificity about cascades in dexterous behavior

The present work is the latest in showing that multifractal formalism can be used to model and discern subtle details within cascade dynamics in task-situated dexterous behavior. The nonlinear interactions weaving events at short and long scales together in cascade dynamics provide an elegant analogy to cyclical, branching relationships between perception and action (e.g. [42]). Nonetheless, this analogy is only so compelling to all stakeholders in the discussion. An unfortunate fact is that single estimates of intermittency, such as a single fractal or a single multifractal-spectrum width, have been opaque. For instance, a single number may govern a scale-invariant cascade, and so a body generating such fractal-like dynamics may be a cascade. However, scale invariance or not, a single number does not articulate the exact pattern and form of relationships [78,80]. So, cascade dynamics has given the impression of being a rebuff or refusal to any discussion of internal structure [79]. In the present work, the spatial and temporal heterogeneity of multifractal-like intermittency across the body has already allowed us to model the role of the internal structure in task satisfaction [52,53]. What the present findings show is that, just as in meteorology and geophysics in which science can model the contributions of sustained streams and currents without presuming to carve firm boundaries between these streams/currents, we too can address the segregation of coordinated behavior into separate cascades. A major strength of multi-scale PDF analysis is its capacity to distinguish between scale-dependent and scale-invariant cascades, and modeling the non-Gaussianity profile with log-timescale using orthogonal polynomials allows us to test hypotheses related to these differently scaling cascades simultaneously.

The present work highlights the importance of bringing together these complementary analytical approaches from the multifractal formalism in future work. This ratcheting up of our potential specificity about internal structure does not entail that we fall back into a program of finding the body part that makes the intermittency. Multifractal geometry resulting from nonlinear interactions across scales can support the development of heterogeneity without founding that heterogeneity in an ontology of strictly independent components that sum together [9,81]. The ontology is sooner of interactions across scale that can prompt the differentiation of functionally specific substreams. Eventually differentiated certainly looks like independence, but we show that this differentiation is sooner a branching from common nonlinear roots than carving at the joints.

Taken together, we present a novel method that allows modeling the task-specificity of physiological fluctuations to address both the spatial form of the PDF and the temporal structure of their sequence. The interactions of multifractal nonlinear temporal correlation (*t*_MF_) with the non-Gaussianity measure obtained via multiscale PDF analysis may offer an elegant blending of disparate uses of the multifractal formalism. This blend may be crucial for the theoretical elaboration of the cascade dynamics underpinning dexterous behavior.

## Supplementary material

**Table S1.** Coefficients of the LME model examining the effects of axis, task, and variable on 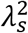 vs. log-timescale curves for interactions with *t*_MF_.

**Table S2.** Coefficients of the generalized linear mixed-effects (GLME) model examining the effects of axis, task, and variable on the growth of lognormality in postural sway with log-timescale.

## Author contributions

D.G.K-S., M.P.F., and M.M. conceived and designed research; D.G.K-S. and M.M. analyzed data; D.G.K-S. and M.M. interpreted results of experiments; M.M. prepared figures; D.G.K-S. drafted manuscript; D.G.K-S., M.P.F., and M.M. edited and revised manuscript; D.G.K-S., M.P.F., and M.M. approved final version of manuscript.

